# Integrated Micropillar Polydimethylsiloxane Accurate CRISPR Detection (IMPACT) System for Rapid Viral DNA Sensing

**DOI:** 10.1101/2020.03.17.994137

**Authors:** Kenneth N. Hass, Mengdi Bao, Qian He, Myeongkee Park, Peiwu Qin, Ke Du

## Abstract

A fully Integrated Micropillar Polydimethylsiloxane Accurate CRISPR Detection (IMPACT) system is developed for viral DNA detection. This powerful system is patterned with high-aspect ratio micropillars to enhance reporter probe binding. After surface modification and probe immobilization, CRISPR Cas12a/crRNA complex is injected into the fully enclosed system. With the presence of double-stranded DNA target, the CRISPR enzyme is activated and non-specifically cleaves the ssDNA reporters initially immobilized on the micropillars. This collateral cleavage releases fluorescence dyes into the assay, and the intensity is linearly proportional to the target DNA concentration ranging from 0.1 to 10 nM. Importantly, this system does not rely on traditional dye-quencher labeled probe thus eliminating the fluorescence background presented in the assay. Furthermore, our one-step detection protocol is performed at isothermal conditions (37°C) without using complicated and time-consuming off-chip probe hybridization and denaturation. This miniaturized and fully packed IMPACT chip demonstrates rapid, sensitive, and simple nucleic acid detection and is an ideal candidate for the next generation molecular diagnostic platform for point-of-care (POC) applications, responding to emerging and deadly pathogen outbreaks.

## Introduction

The widespread impact of the current coronavirus (COVID-19) is a striking indicator of the fact that the global community is struggling to battle infectious diseases. Failure to contain the virus early has resulted in an outbreak that has infected over 120,000 people with over 3,000 deaths.^1,2^ The USA CDC predicts that a nationwide epidemic is unavoidable.^3^ The current situation highlights a urgent need for access to real-time detection that is accurate and effective to identify those infected so they can be properly quarantined and treated.^4,5^ An effective vaccine could be the best solution to contain epidemics. However, vaccines take a very long time to develop, as evidenced with the African swine fever virus (ASFV), which was initially discovered in 2018 but only had a preliminary vaccine announced in January of this year.^6^ It took over an additional month to show its effectiveness in laboratory tests, and it still needs to be proven effective in the field.^7^

Currently, there are no proven treatments for either COVID-19 or ASFV. Even if one were to be developed soon, it can take years to prove its effectiveness in clinical trials and mass produce the vaccine, not to mention distribute and administer it in affected areas.^8^ To contain and prevent the spread of these contagious outbreaks, a rapid point-of-care (POC) testing device is essential. Current methods can take up to two days for tittering in a centralized laboratory in order to diagnose whether a sample contains the disease, and is not viable and inefficient when trying to isolate those infected and prevent them from spreading the disease to others.^9,10^

In addition, the collected patient samples need to be sent to a centralized laboratory, leading to very long turnaround times and greatly limiting the number of people who could be tested and confirmed.^11^ Strides have been made to develop POC detection methods which can be deployed in the field, and currently lab-on-chip (LOC) devices or simple test kits which involve real-time polymerase chain reaction (PCR) have emerged as one of the leading choices for meeting the desired criteria.^12^ PCR is particularly sought after due to its ability to amplify the viral RNA/DNA from a few copies to billions.^13^ It has also been shown to work with clinical samples in LOC devices with low volumes of both reagents and patient samples.^14^ However, there have been many issues with implementing PCR test kits, such as a shortages in test kits and trained professional to use them, as well as potential inaccurate results from the kits.^15,16^

Clustered regularly interspaced short palindromic repeats (CRISPR) provides an alternative to PCR amplification techniques for the detection of viral RNA/DNA. Certain Cas proteins, such as CRISPR-Cas12a, have been shown to be powerful in biological detection due to their ability to indiscriminately cut single-stranded DNA (ssDNA) after they are activated by a target DNA.^17^ This is extremely useful when paired with “reporter probes” (ssDNA strands with a fluorescent dye and quencher attached to them), as the CRISPR complex can cleave the reporter probe and release the dye for fluorescence quantification. However, one of the issues with utilizing the CRISPR complex is the high fluorescence background signal associated in the sample as the quencher cannot fully quench the fluorescent dye.

Solid-phase detection assays have been developed as one potential solution to overcome this issue presented in the liquid phase. On the solid surface, reporter probes do not require a quencher since they are only measured in the liquid phase after degradation, thus no fluorescent signal will be detected without the target DNA present in the assay. However, an extended surface with a larger surface area is always needed to increase the probe binding capacity. It has been shown that these extended surface can increase the amount of probe binding, lower the detection limit of the target of interest, and extend the detection dynamic range.^18,19^

Here, we present a fully enclosed Integrated Micropillar Polydimethylsiloxane Accurate CRISPR Detection (IMPACT) system for nucleic acid target detection. The reporter probes were firstly immobilized in the enclosed channel and the CRISPR complex was pumped into the system for reaction (Fig. 1a). Leveraging the high activity of CRISPR-Cas12a enzyme and the ability of micropillars to bind more reporter probes, we successfully detect double-stranded DNA target without background issues. In addition, the IMPACT chip requires low volumes of reagents, operates in a one-step detection fashion, and does not require complicated temperature control, making it ideal for POC applications. To prove the concept of the IMPACT chip for viral DNA detection, the target sequence selected for the crRNA was synthesized after a segment of the ASFV-SY18 genome (B646L). In the future, our device could work with any RNA or DNA based pathogens, such as the emerging COVID-19.

**Figure 1:**
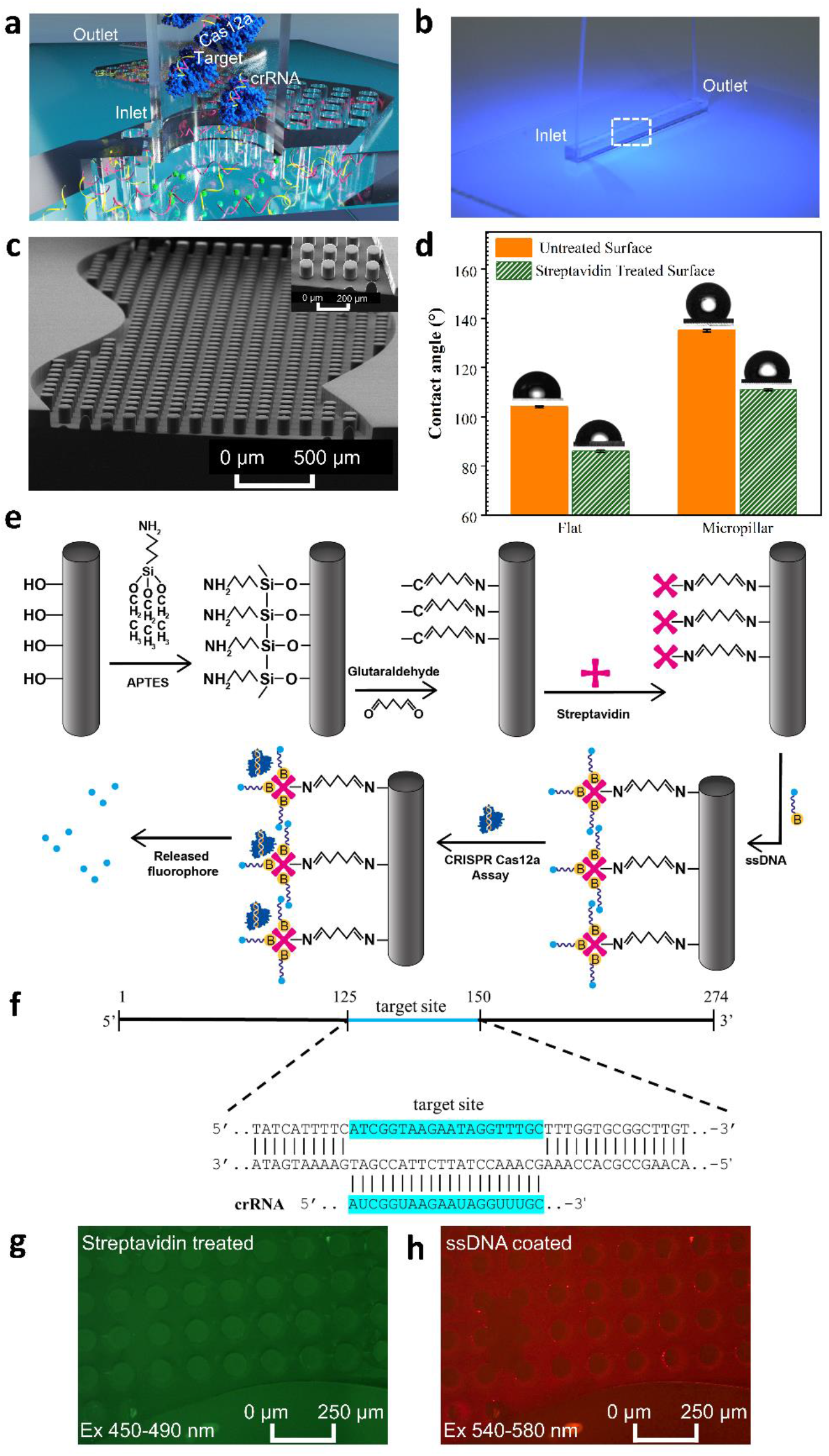
a) Schematic of the CRISPR based IMPACT chip DNA detection. b) Photograph of the IMPACT Chip. The dashed white box indicates regions patterned with micropillars. c) SEM image of the micropillars. Inset: high magnification image. d) Static water contact angle measurement on both flat and micropillar surface before (orange) and after (green) surface treatment. e) Schematic of the surface treatment protocol, ssDNA reporter probe binding, and CRISPR detection. f) ASFV target DNA sequence and the corresponding crRNA sequence. g) Fluorescent image of channel which received chemical treatment and streptavidin binding (green fluorescence). h) Fluorescent reporter probe covered IMPACT chip, showing red fluorescence (emission: 668 nm).

## Results

The IMPACT chip (Fig. 1b) consists of 6 cm long channels patterned with Polydimethylsiloxane (PDMS) micropillars. Fig. 1c shows the SEM image of the channel, revealing periodic curved nature of the channel and even spacing of the micropillars (diameter: 100 μm; height: 120 μm). After channel fabrication and sealing, surface treatment with APTES (3-Aminopropyl)triethoxysilane) and glutaraldehyde was performed for streptavidin immobilization.^20^ After streptavidin coating, the static water contact angle decreases from ~105 degree to ~90 degree for a flat PDMS surface and ~138 degree to 110 degree for micropillar surface, respectively (Fig. 1d). Fig. 1e shows the entire sample preparation and detection protocol. DNA reporter probes with a biotin label were conjugated on the streptavidin coated micropillars. CRISPR Cas12a complex was then introduced into the channel. With the presence of the ASFV target DNA, it cleaves the reporters from the micropillars for detection. The sequence of the DNA target and the crRNA are listed in Fig. 1f. The target was selected based on the PAM region in the ASFV-SY18 genome.^21^ It has been shown that auto-fluorescence occurs from the chemical treatment of APTES and glutaraldehyde.^22^ In our case, we were able to confirm the phenomenon was specific to glutaraldehyde and APTES (Fig. SI 1a), and use it to show that it only occurs on PDMS in the presence of these chemicals (Fig. SI 1b). Fig. 1g shows the uniformity of the chemical treatment on the surface, as the intensity of the auto-fluorescence (emission: 525 nm) is uniform throughout. We then incubated the reporter probes (emission: 668 nm) in the channel for 3 hrs and showed uniform coating in the channel (Fig. 1h).

After establishing a surface modification and streptavidin incubation strategy, we first determined the channel washing conditions. After incubating biotinylated photocleavable capture probes (/5PCBio/TTATTCTTATTGTGTGAACTGCTCCTTCTTGACTCCACC/36-FAM/) in the channel for 3 hrs, the channel was washed with 75 μL DI water. Then, the channel was evacuated, and the fluorescence intensity of the supernatant was immediately evaluated. After the first wash, a high fluorescence peak was observed for all the samples (inset of Fig. 2a), indicating that excessive DNA probes were washed from the channel. A second wash was performed to confirm the unbounded DNA probes were completely removed from the channel. After the second wash, the collected supernatant barely shows any fluorescence peak. As shown in Fig. 2a, the integrated signal (490 to 700 nm) for the first supernatant ranges between ~8,000 to 9,000 counts, regardless of surface treatment conditions (Saturated the spectrometer due to high signal). On the other hand, the integrated signal for the second supernatant is only ~300 counts, eliminating the background caused by weakly bounded DNA probes. After washing, the bounded reporter probe was released by UV exposure and the results are presented in Fig. 2b. The flat channel coated with streptavidin shows ~3 fold more reporter probe binding than the uncoated surface. In addition, the micropillar channel coated with streptavidin shows the highest signal among the three substrates.

**Figure 2:**
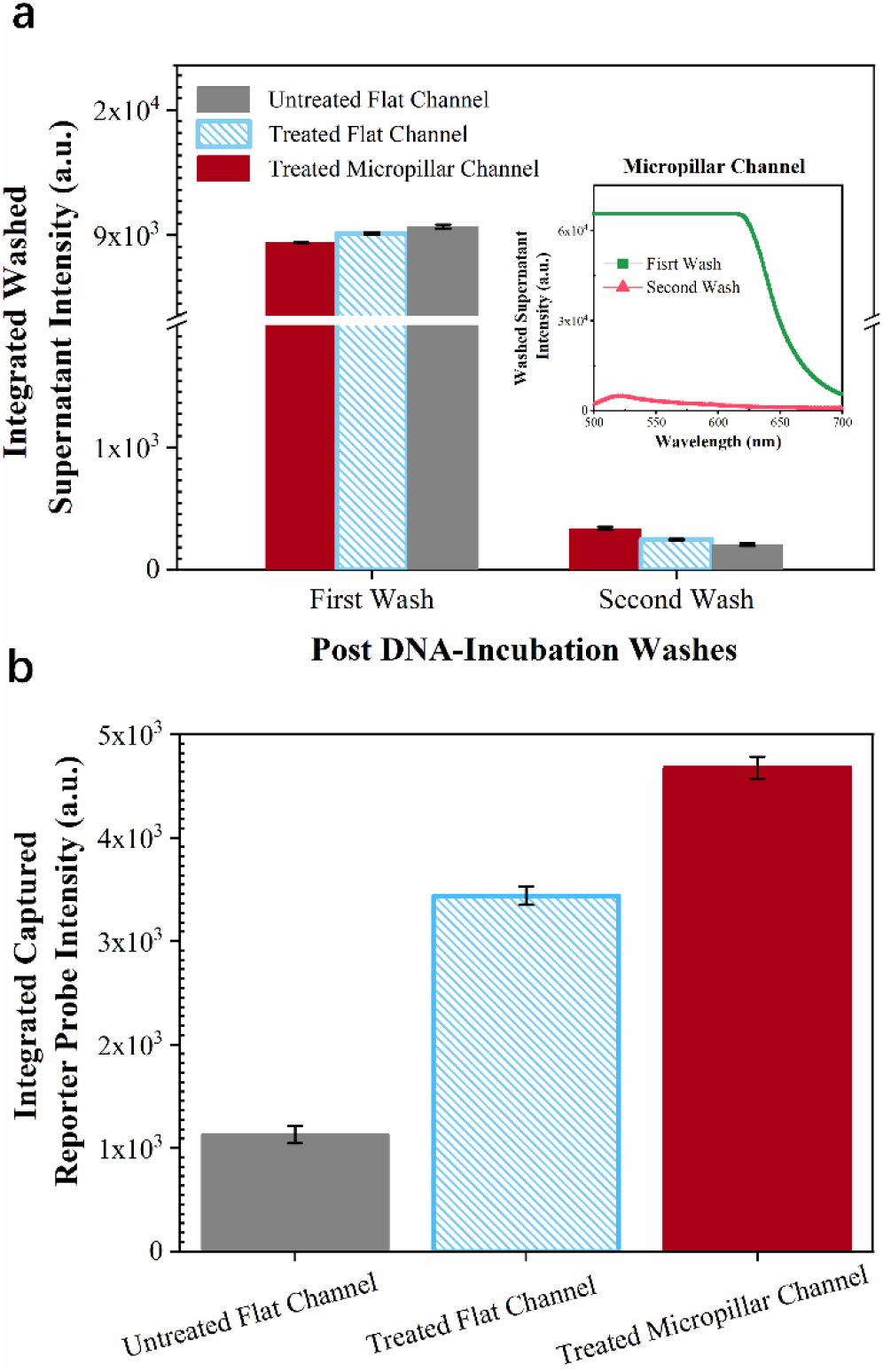
Washing Data for flat and micropillar channels. The treated flat and microchannel samples were coated with streptavidin. (a) Integrated intensity of the supernatant after washing with 75 uL of DI water through the channel. Error bars are standard deviation of the mean. Inset shows uncorrected emission curve of washed reporter probe for the treated micropillar channel. (b) Integrated intensity of released reporter probes with UV light exposure.

With an optimal washing condition in hand, we then studied the effect of incubation time for biotin labeled DNA binding on streptavidin coated PDMS surface. After channel surface modification and streptavidin treatment, fluorescent reporter probe was introduced to bind on the solid surface. Then, the washing protocol was performed, followed by UV light exposure to retrieve the reporter probe. As shown in Fig. 3, for streptavidin coated surface, the number of DNA immobilized on the surface does not show significant change with an incubation time between 10 to 60 min as the integrated fluorescence intensity of the retrieved DNA ranges between 50,000 to 60,000 counts. However, the binding capacity increases ~25% with 3 hrs incubation when compared to 10 min incubation. Although the binding capacity further increases with 24 hrs incubation, it requires refrigeration as the streptavidin protein could denature at room temperature. On the other hand, the negative sample without streptavidin coating does not show a correlation to the incubation time, indicating less specific binding. Moreover, the negative sample always shows much lower fluorescence intensity than the positive sample. For example, with 3 hrs incubation, the retrieved DNA from positive sample is ~2.6 times higher than the negative sample. It further indicates that surface modification can enhance the probe binding capacity. Therefore, we used 3 hrs incubation to prepare the IMPACT chip for solid phase CRISPR detection.

**Figure 3:**
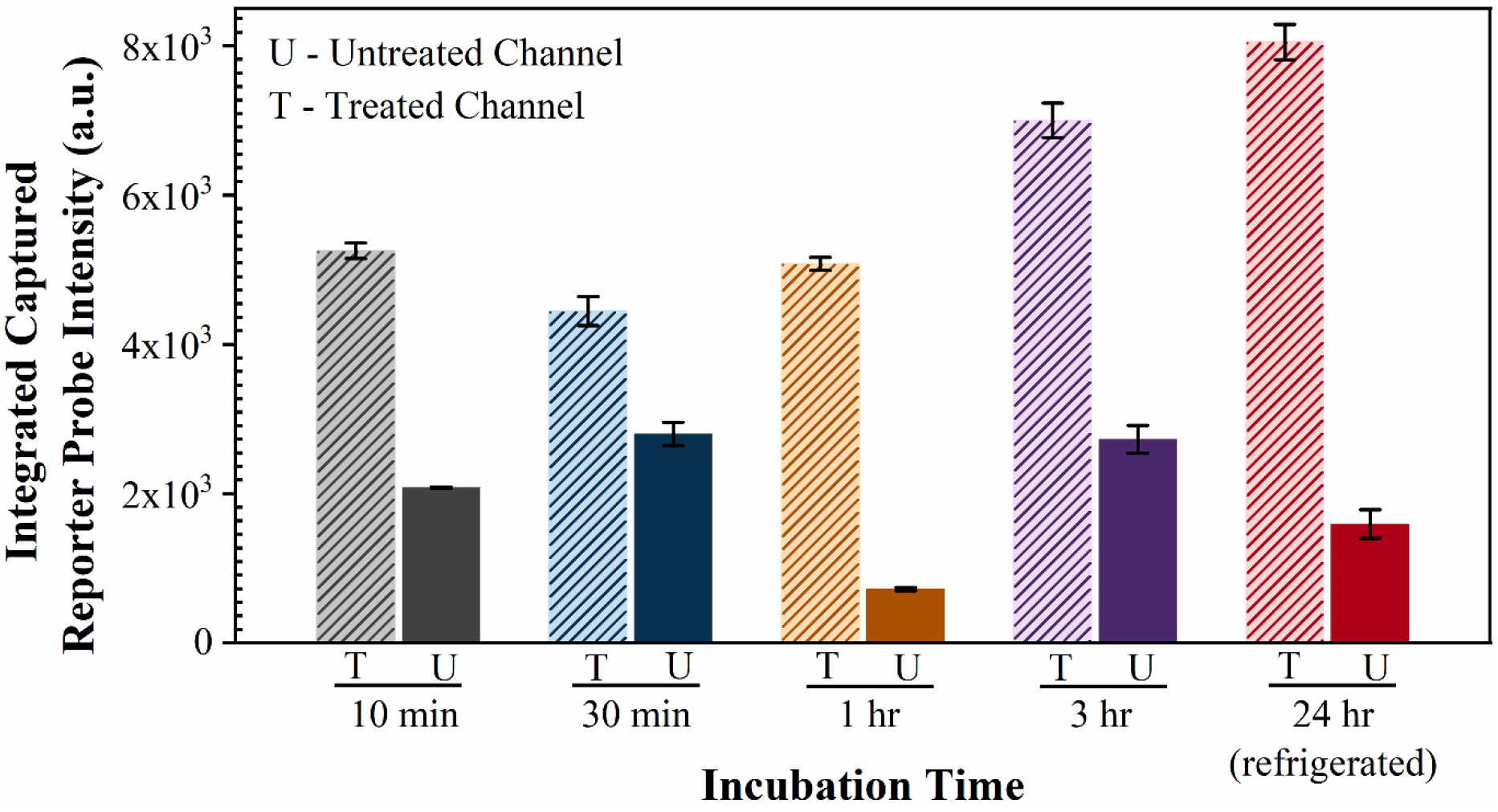
Released reporter probe intensity versus incubation time (10 min to 24 hrs). Treated samples received surface modification and streptavidin treatment before photocleavable reporter probe incubation (1×10^6^ nmoles). The sample was retrieved via UV photocleavage. Error bars are standard deviation of the mean

The extended surface provided by high-aspect ratio micropillars significantly increases the reporter probe binding capacity. To demonstrate this, we compared the number of captured DNA molecules on the micropillar channel with the flat channel (Fig. 4). With an input of 1×10^4^ nmoles, the retrieved DNA from the micropillar channel has an integrated intensity of ~2,000 counts, which is much higher than the flat surface (~900 counts). With an input of 1×10^5^ nmoles, the retrieved DNA from the micropillar channel has an integrated intensity of ~5,500 counts and is ~40% higher than the flat channel. With an input of 1×10^6^ nmoles, the micropillar channel still shows higher signal than the flat channel but is comparable to 1×10^5^ nmoles sample, indicating that the channel is saturated with an input of ~1×10^−5^ nmoles. Thus, we used 1×10^5^ nmoles reporter probes in our CRISPR detection as higher load could cause unnecessary background.

**Figure 4:**
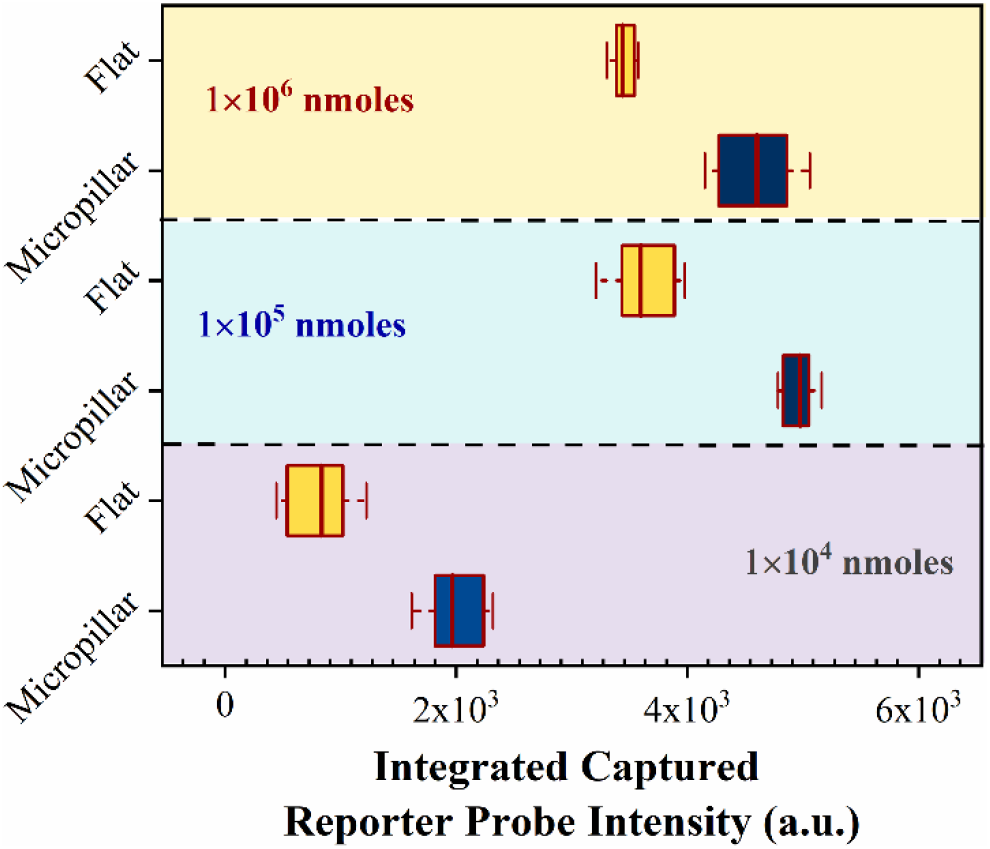
Released ssDNA reporter probe integrated intensity for flat and micropillar channels with a DNA input load of 1×10^4^, 1×10^5^, and 1×10^6^ nmoles, respectively. UV photocleavage was used to retrieve the ssDNA reporter probes. Error bars represent 5%-95% confidence intervals and middle line is mean.

We combined the DNA probe modified micropillar channel with CRISPR Cas12a assay for solid-phase and background free viral DNA sensing. We first mixed CRISPR Cas12a/crRNA/target DNA in an Eppendorf tube and then injected the complex into the IMPACT chip and allowed it to incubate for 2 hrs. The activated complex diffuses in the microchannel and non-specifically cleaves the reporter probes from the micropillars. The uncorrected emission curve and the integrated fluorescence signal of the CRISPR experiments are shown in Fig. 5a and 5b, respectively. The measured fluorescence intensity linearly increases with the target concentration ranging from 0.1-10 nM (Pearson’s R=0.9653). On the other hand, the supernatant without any ASFV target DNA input does not present a fluorescence signal as the CRISPR Cas12a cannot be activated, demonstrating a fully enclosed and efficient microdevice for background-free viral DNA quantification.

**Figure 5:**
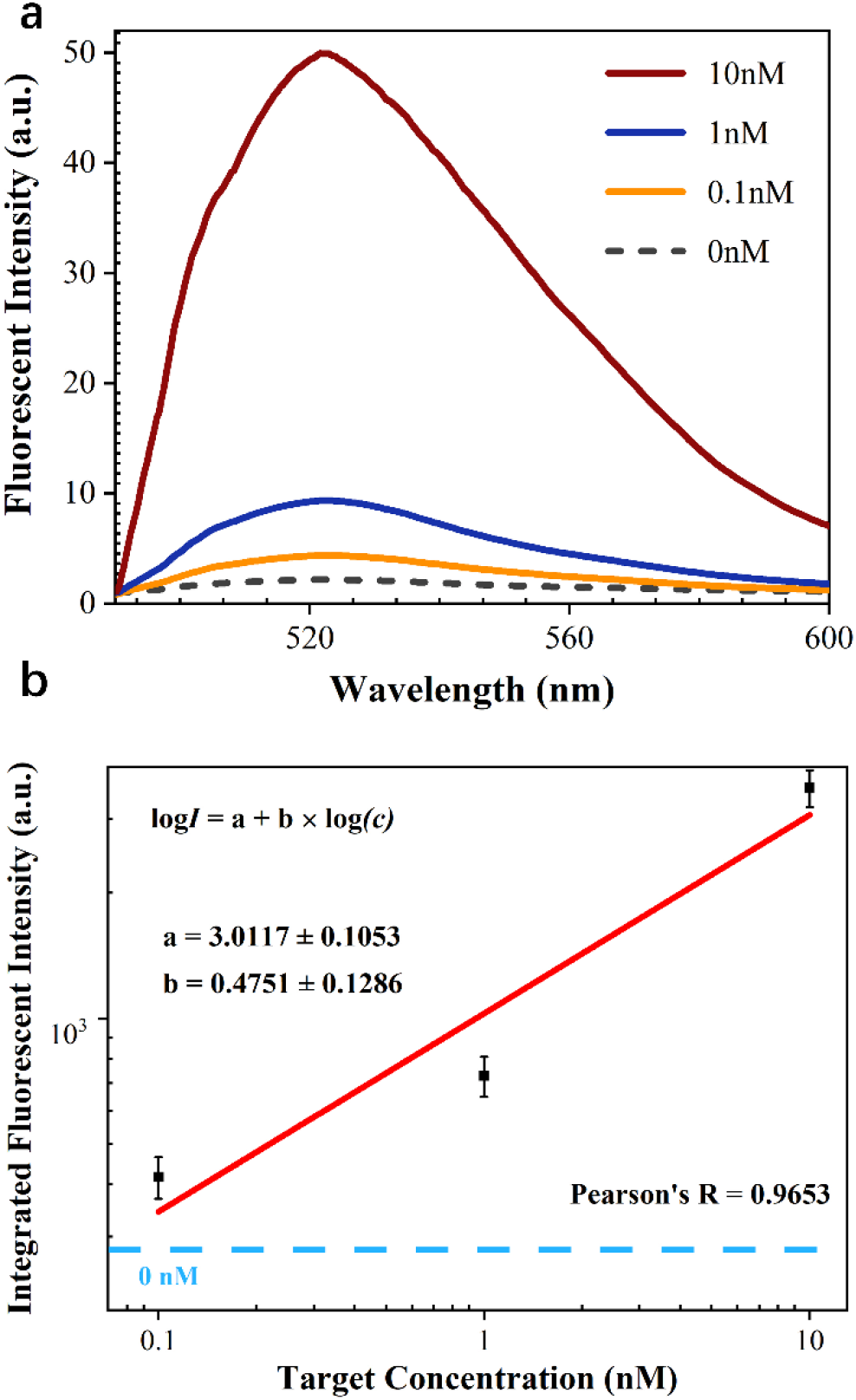
IMPACT chip performance with different ASFV target DNA concentrations. a) Uncorrected emission curve of the collected supernatant with a target concentration ranging from 0.1 nM to 10 nM. b) Integrated intensity of the collected supernatant versus target concentration on a logarithmic scale (Pearson’s R=0.9653). Error bars are standard deviation of the mean.

## Discussion

We have shown that the PDMS micropillars provide a significant increase in solid-state capture probe binding than the flat surface. The increased ssDNA capture probe capacity can extend the detection limit and also dynamic range.^23,24^ In addition, the micropillar array increases the interaction between biomolecules and the substrate, as the surface area in contact with any given solution is significantly higher. To further increase the probe binding capacity, we can increase the aspect ratio of PDMS micropillars and the microchannel length. Indeed, the channel was designed to have a periodic curvature throughout its length in order to show the ability to utilize a curved design with the chip, which in the future could snake back and forth across an entire sample, greatly increasingly the overall surface area. The aspect ratio of the microstructures can be further increased by choosing more rigid materials. For example, an aspect ratio of 160:1 has been demonstrated on silicon by using deep-ion reactive etching.^25,26^ Silicon would also have the advantage of potentially removing some of the air bubbles seen during use with the IMPACT chip, which could ensure greater uniformity in coating of reporter probes. Alternatively, the emerging additive manufacturing technology can also be used to create such ultra-high aspect ratio microstructures as an efficient target capture platform.^27,28^

One of the main advantages of the IMPACT chip compared to traditional CRISPR assays is its ability to limit the background caused by dye-quencher probes, which is typically seen in CRISPR detection in the liquid state and needs to be designed around to lower the detection limit.^29,30^ Our device utilizing solid-phase CRISPR does not need to tether a quencher on the probe. The cleaved CRISPR products are sent to a separate reservoir for detection thus completely avoiding the fluorescence background caused by the tradition “one-pot” detection. As shown in Fig. 5, without the presence of ASFV target DNA, no fluorescence background signal was detected, demonstrating this powerful background-free detection. This is an important improvement for molecular diagnostics, especially in low light settings in which the amount of background fluorescence present can significantly affect the detection limit.^31,32^ Leveraging this advantage with increasing the aspect ratio either with PDMS or a silicon substrate will be able to further increase the accuracy and robustness of the device and improve the limit of detection.

The IMPACT chip offers a simple one-step detection strategy and does not require off-chip incubation. Traditional nucleic acid based detection requires stringent multi-probe hybridization and washing thus is slow and complicated.^33,34^ In this work, the CRISPR complex is quickly mixed with the target sample and then the CRISPR/crRNA/target DNA complex is injected into the channel for on-chip detection. The chip sits on a hotplate set at 37°C and does not need thermal cycling with temperature control like PCR does. ^35,36^ Thus, the detection system we developed is much simpler and more compact, ideal for POC applications.^37,38^ To extend the detection limit, our system can be integrated with isothermal amplification methods such as recombinase polymerase amplification for sensitive “one pot” target amplification and CRISPR reaction, without the background issues seen in traditional “one-pot” methods.^17,39^

An advantage of the IMPACT chip compared to other detection apparatuses such as the SHERLOCK test strip is that it is a fully enclosed system without extra packaging need.^40,41^ This is advantageous as the treated micropillars are sealed before molecular diagnostics. The reporter probes are immobilized within the channel, avoiding degradation issues from outside contaminants. This is particularly important when dealing with RNA target as it is susceptible to degradation by the presence of RNase.^42^ Our fully integrated chip without special packaging needs is able to avoid RNase exposed in air, dust, and human hands in POC settings.

This IMPACT chip concept can be extended to detect many biomarkers such as exosomes^43^, single cells^44^, and surface proteins^45^. For example, the detection sensitivity of traditional immunoassay approaches is limited due to the low binding capacity. Magnetic beads have been widely used to increase the binding affinity.^46,47^ However, it is difficult to incorporate beads into the microfluidic channel as they easily settle down and get stuck in the channel. In vivo detection is also challenging as the beads emit strong auto-fluorescence. The micropillar arrays that we developed are patterned in the chip and can avoid those problems. In the future, on-chip treatment of blood infections can be achieved by immobilizing proper capture probes onto the IMPACT chip.^48,49^

## Materials and Methods

### Device Fabrication

To create the IMPACT microfluidic chip, a mold was created using standard photolithography on a 100 mm silicon wafer. The silicon wafer was first dehydrated by baking it at 200°C for 20 min on a hot plate. After cooling to room temperature, SU-8 2075 (Microchem) was spun coat onto the silicon wafer with a thickness of 120 μm. The wafer was then left to sit for 10 min on a level surface to allow for reflow, and then soft baked on a hotplate at 65°C for 10 min, followed by 20 min at 95°C. After that, the wafer was exposed using a Karl Suss MA6 Mask Aligner for 15-20 s. It was then allowed to sit for 5 min, before receiving a post exposure bake at 80°C for 10 min. The sample was developed for ~10 min in SU-8 developer (MicroChem) and then sprayed down with IPA and dried. A final hard bake at 145°C was done on a hotplate for 5 min. After the hard bake, the wafer was silanized overnight using Silanization Solution I (~5% dimethyldichlorosilane in heptane) in a desiccator.

The PDMS channel was created by pouring 10:1 ratio of SYLGARD 184 Silicone Elastomer Base to SYLGARD 184 Silicone Curing Agent over the SU-8 molds and allowing it to cure. The PDMS was then peeled from the SU-8 mold, and holes (diameter of 1 mm) were punched at both ends of the channels to allow for flow of reagents through the channel. The channels were then cleaned in an ultrasonic bath for 5 min using ethanol, dried, and then cleaned again in the same manner using deionized water. The PDMS slab was then bonded to a pre-cleaned glass substrate by treating both with oxygen plasma (Electro-technic products) for 2 min, followed by pressing the substrate and PDMS together. The device was immediately baked on a hotplate overnight at ~125°C.

### Surface Modification

The device was filled with 10% APTES (Sigma Aldrich, (3-Aminopropyl)triethoxysilane) in Ethanol (BVV, Lab Grade Ethanol 200 Proof USA Made). A total of 75 μl of the solution was flown into the channels at a flow rate of 15 μl/min using a syringe pump (WPI, SP2201). The APTES solution was then left to incubate in the channel for 10 min and washed with 96% ethanol. After washing, the remaining liquid in the channel was drained by a syringe filled with air, and then with a compressed air canister (Dust Off, Electronics compressed-gas Duster) to dry the channel. The channel was then baked on a hotplate at 125°C for 30 min. Glutaraldehyde (AD) solution (25%) in DI water was next flown to fill the channel in the same manner as described above, but was allowed to incubate for a full hour after the 75 μl of solution was flown through. The channel was then flushed with DI water and dried with air.

### Streptavidin Immobilization

After surface treatment, each channel was injected with 20 μl of Streptavidin (ThermoFisher Scientific, Streptavidin S888) at a concentration of 4 mg/ml using a 23 GA microliter syringe (Hamilton). Since each channel has a volume of ~14 μl, any excess solution was allowed to exit to the outlet. The solution was left in the channel for 2 hrs to incubate.

### Surface Characterization

A goniometer system (Model 260, Ramé-hart) was used to characterize the chemically treated PDMS surface. A sessile droplet (~4 μL) of deionized water was placed on the PDMS surface with an automated dispensing system (Part No. 100-22). The static water contact angle was immediately measured by using the DROPimage Matrix software, provided by Ramé-hart.

### Reporter Probe Immobilization

In a similar manner as the streptavidin was applied, 20 μl probe solution in PBS buffer (Gibco TM, PH 7.4) was injected into the channel with a microliter syringe and allowed to incubate for 3 hrs, regardless of whether a photocleavable linker was attached to DNA bases.

### Ultraviolet (UV) Cleavage

Reporter probe with a UV cleavable linker was cleaved by using a UV light (wavelength: 311 nm). Briefly, the sample was placed under a UV light with the glass side of the channel facing the light, at a distance of ~5 cm from the UV lamp to the glass sealing the channel. The light was then turned on and the sample was exposed to the UV light for 10 min. Finally, the released reporters were retrieved by injecting 75 ul of DI water into the channel.

### Solid-Phase CRISPR Cas12a Detection

LbCas12a (New England BioLabs, Inc.) with a concentration of 50 nM was pre-assembled with 62.5 nM crRNA (IDT, Inc.) at room temperature for 10 min. Then, LbCas12a-crRNA complexes were mixed with 1 x Binding buffer and 14.75 μl Nuclease Free Water (IDT, Inc.) to reach a 20 ul reaction volume. After adding ASFV target DNA with various concentrations into the mixture, the detection assay was activated and immediately injected into the IMPACT chip. The device was incubated on a hotplate at 37°C for 2 hrs to allow for optimized reaction. Finally, the cleaved product was retrieved from the channel and evaluated by a custom designed fluorometer.

### Fluorescence Quantification

To quantify the fluorescence intensity of the ssDNA reporter probes, a custom designed fluorometer was used and has been reported before.^21,50^ Briefly, a continuous wave laser with an emission peak at 488 nm (Sapphire 488 LP, Coherent) was aligned under a reservoir filled with fluorescent molecules. The fluorescence signal was collected by an off-axis parabolic mirror and a fiber coupled mini USB spectrometer (USB 2000+, Ocean Optics). To reduce background noise from the excitation light, a 488 nm notch filter (Thorlabs, Inc.) was placed in front of the optical fiber.

## ASSOCIATED CONTENT

Fluorescence images of the surface modified with APTES - Glutaraldehyde (positive control) and APTES-Valeraldehyde (negative control). Negative control images of Figure 1g and 1h which did not receive any surface modification.

## Author Contributions

K.H., M.B., and K.D. designed the experiments. K.H., M.B., and Q.H. conducted the experiments. K.H., M.B., and K.D. wrote the manuscript. All the authors commented on the manuscript.

‡These authors contributed equally. The manuscript was written through contributions of all authors.

## ACKNOWLEDGMENT

This research used resources of the Center for Functional Nanomaterials, which is a U.S. DOE Office of Science Facility, at Brookhaven National Laboratory under Contract No. DE-SC0012704. The authors would like to thank Wenrong He and Personalize Healthcare Technology (PHT180) at the RIT for schematic design.

## ABBREVIATIONS

IMPACT: Integrated Micropillar Polydimethylsiloxane Accurate CRISPR Detection
ASFV: African Swine Fever Virus
POC: Point-of-care
LOC: Lab-on-chip
PCR: Polymerase chain reaction
CRISPR: Clustered regularly interspaced short palindromic repeats
ssDNA: Single-stranded DNA
PDMS: Polydimethylsiloxane
APTES: (3-Aminopropyl)triethoxysilane)

## TOC only

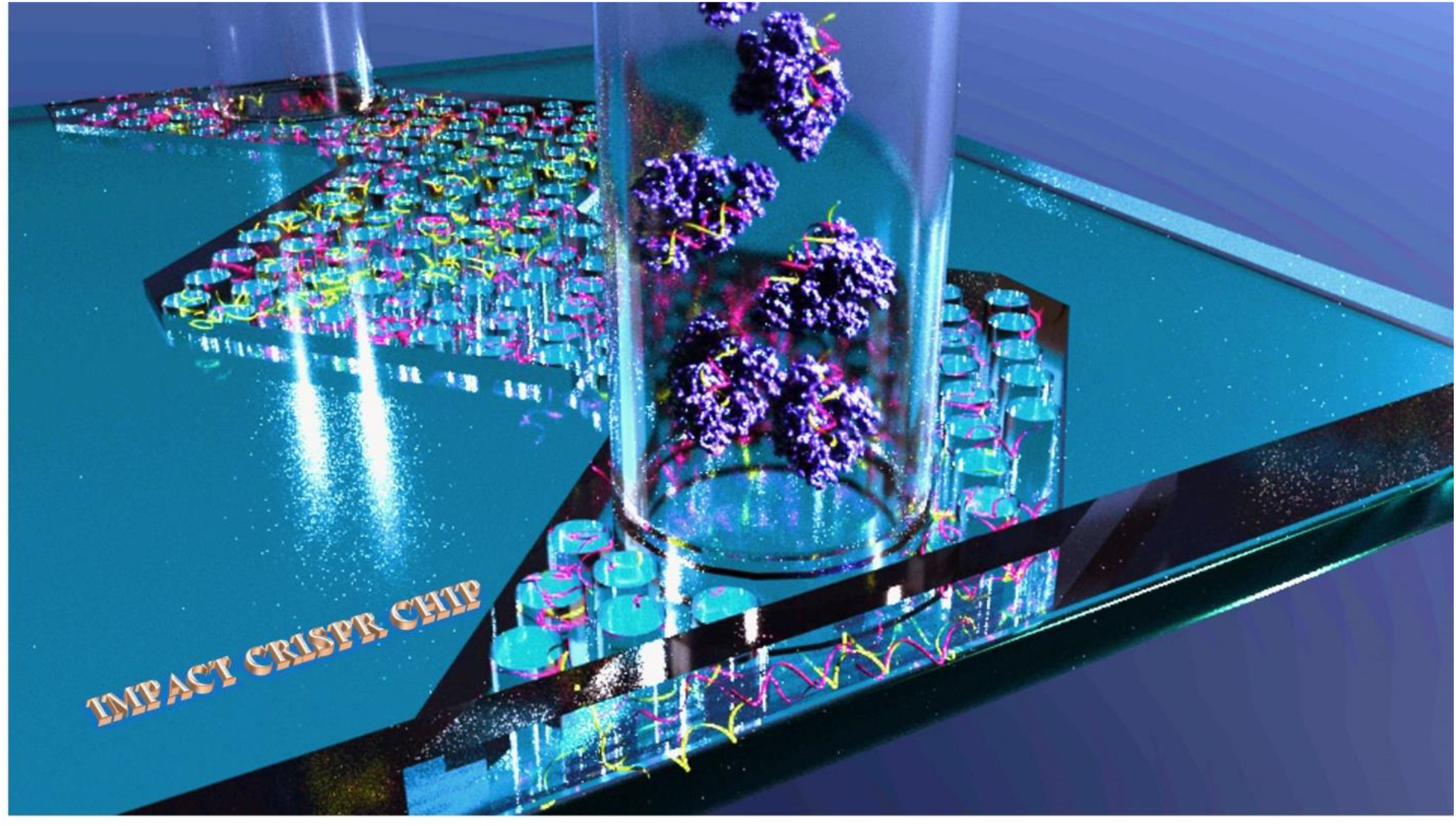

